# Neural dynamics encoding risky choices during deliberation reveal separate choice subspaces

**DOI:** 10.1101/2025.04.29.651314

**Authors:** Logan M. Peters, Alec Roadarmel, Jacqueline A. Overton, Matthew P. Stickle, Jack J. Lin, Zhaodon Kong, Ignacio Saez, Karen Anne Moxon

## Abstract

Human decision-making involves the coordinated activity of multiple brain areas, acting in concert, to enable humans to make choices. Most decisions are carried out under conditions of uncertainty, where the desired outcome may not be achieved if the wrong decision is made. In these cases, humans deliberate before making a choice. The neural dynamics underlying deliberation are unknown and intracranial recordings in clinical settings present a unique opportunity to record high temporal resolution electrophysiological data from many (hundreds) brain locations during behavior. Combined with dynamic systems modeling, these allow identification of latent brain states that describe the neural dynamics during decision-making, providing insight into these neural dynamics and computations. Results show that the neural dynamics underlying risky decision, but not decisions without risk, converge to separate subspaces depending on the subject’s preferred choice and that the degree of overlap between these subspaces declines as choice approaches, suggesting a network level representation of evidence accumulation. These results bridge the gap between regression analyses and data driven models of latent states and suggest that during risky decisions, deliberation and evidence accumulation toward a final decision are represented by the same neural dynamics, providing novel insights into the neural computations underlying human choice.

**Highlights:**

- Highly accurate decoding can be accomplished with dynamical systems modeling that revealed distinct attractor-like subspaces, one for each of the two options (gamble or safe bet).
- Early during deliberation, single trial neural trajectories show rapid transitioning between these subspaces, but as the time of choice selection approached, the single trial trajectories converged towards one of the subspaces, whose identity was consistent with the subject’s choice.
- These dynamics are specific to risky decisions as we found classification accuracy dropped to chance for trials where the gamble option was guaranteed to be successful (100% win probability) or unsuccessful (0% win probability).
- These results suggest that deliberation and evidence accumulation toward a final decision in the presence of any risk can be represented by the same neural dynamics, providing novel insights into the neural computations underlying human choice.

## 1. Introduction

Human decision-making involves the coordinated activity of multiple brain areas, acting in concert, to enable humans and animals to adaptively make choices (Ebitz et al., 2020; Li et al., 2013; O’Connell et al., 2018; X. J. Wang, 2012). Increasing evidence (Kennerley et al., 2011; Wallis & Miller, 2003)from animal models supports a distributed view of neural activity underlying reward-guided behavior, with activation across brain regions emerging simultaneously rather than sequentially during the deliberation period prior to choice selection (Kennerley et al., 2011; Man et al., 2024; Ottenheimer et al., 2023; Steinmetz et al., 2019; Wallis & Miller, 2003) . Animal studies have shown that localized neural activity in a variety of regions reflects specific reward-related computations such as risk (Preuschoff et al., 2006; Schultz et al., 1997), value (Enel et al., 2020; Gottfried et al., 2003; Hayden & Niv, 2021; Kennerley et al., 2011; Kobayashi et al., 2021; Padoa-Schioppa & Assad, 2006, 2008; Paton et al., 2006; Polanía et al., 2014; Rangel et al., 2008; Smith et al., 2010; Strait et al., 2015; Usher et al., 2012; Wallis, 2012; Williams et al., 2021)and reward probability (Akaishi & Hayden, 2016; Balleine et al., 2007; Bayer & Glimcher, 2005; Farashahi et al., 2019; Gauthier & Tank, 2018; Gottfried et al., 2003; Gruber et al., 2013; Hayden et al., 2009; Hira et al., 2014; Hunt & Hayden, 2017; Kishida et al., 2016; Man et al., 2024; Saez et al., 2018; Schultz, 2017; Schultz et al., 1997, 2008; Sosa & Giocomo, 2021; Tremblay & Schultz, 1999; M. Z. Wang & Hayden, 2017). However, how this sub-second neural activity during the deliberation period enables choice, including the role of neural activity across different frequency bands and brain regions, is not well understood in the human brain, partially due to the difficulty of recording neural activity in multiple brain areas.

To directly observe human neural activity with high anatomical precision, signal-to-noise and temporal resolution we leveraged human intracranial electroencephalography (iEEG) recordings from neurosurgical patients. We combined these recordings with reward tasks to study the relationship between distributed neurophysiological activity and choice behavior. This approach overcomes some of the limitations of non-invasive human neural methodologies including limited signal-to-noise ratio (EEG), temporal resolution (fMRI), or anatomical accuracy (EEG) (Buzsáki et al., 2012). Patients played an economic risky decision-making game while local field potentials (LFP) were recorded from multiple reward-related brain areas, including orbitofrontal, lateral prefrontal, premotor and motor cortices, insula, and amygdala (Saez et al., 2018). The game included trials where there was no risk (the outcome was certain) as well as trials with varying probabilities of resulting in a win (win probability).

To examine distributed neural dynamics during deliberation, we developed models that predicted single trial behavior (choice) and used them to assess how the information about choice is conveyed by changes in power across frequency bands and regions. Our models applied dimension reduction followed by classification techniques and were used first to identify the features that conveyed the most information about choice by evaluating the classifier performance. We then compared performance across models to identify candidate neural mechanisms that might be used by the brain to compute the subject’s final choice. Finally, we examined the output of the models to gain insight into the relationship between changes in neural activity and behavior. As a first approximation, we examined the linear representation of neural computations underlying choice using two approaches: Principal Components Analysis (PCA) that describes how the covariance of data at different frequencies across different brain regions combine to encode choice and Linear Dynamical Systems (LDS) modeling that captures the linear component of how the neural activity at one moment in time produces the activity at the next (Cheng & Sabes, 2006; Tóth, 2010; Yu et al., 2008). While there are likely critical non-linear computations, the aggregate activity of large populations of neurons reflected by the on-going oscillatory activity is likely to have a substantial linear component (Rich & Wallis, 2016) and an understanding of this linear component will provide both insight into how, when and where non-linear computation are important, as well as a baseline for how much information can be conveyed linearly.

Our results show that information about choice can be represented in a relatively low dimensional state-space defined seven Principal Components (PCs) or seven latent variables (LVs) derived from the LDS model using high-frequency activity[30-200Hz]. Highly accurate decoding can be accomplished with dynamical systems modeling that revealed convergence to two subspaces, one for each of the two options (gamble or safe bet). Early during deliberation, single trial neural trajectories show rapid transitioning between these subspaces, but as the time of choice selection approached, the single trial trajectories converged towards one of the subspaces, whose identity was consistent with the subject’s choice. These dynamics are specific to risky decisions as we found classification accuracy dropped to chance for trials where the gamble option was guaranteed to be successful (100% win probability) or unsuccessful (0% win probability). These results suggest that deliberation and evidence accumulation toward a final decision in the presence of any risk can be represented by the same neural dynamics, providing novel insights into the neural computations underlying human choice.

## 2. Methods

### 2.1 Subjects

Data were collected from 34 subjects (19 female) with intractable epilepsy who were implanted with chronic subdural grid or strip electrodes (electrocorticography, ECoG) or stereotactic EEG (sEEG) electrodes as part of a procedure to localize the epileptogenic focus. Electrode placement was based solely on the clinical needs of each subject. Fourteen subjects had psychometric curves outside of the range recorded from healthy subjects ((Saez et al., 2018)), suggesting either they didn’t understand the game or were not paying attention during the game, and were, therefore, excluded from further analysis (Supplemental Data Figure 5). For the remaining 20 subjects, we excluded electrodes that showed evidence of epileptic activity. Data were recorded postoperatively in the epilepsy monitoring unit at five hospitals: The University of California (UC), San Francisco Hospital (n = 3), the Stanford School of Medicine (n = 3), UC Irvine Medical Center (n = 23), and UC Davis Medical Center (n = 5). As part of the clinical observation procedure, subjects were off anti-epileptic medication during these experiments. All participants gave written informed consent to participate in the study in accordance with the University of California, Davis or University of California, Berkeley Institutional Review Board. Subjects understood that they could decline participation at any time, and verbal assent was reaffirmed prior to each experimental task.

### 2.2 Electrophysiological data acquisition

Subjects underwent either electrocorticography (ECoG), providing subdural coverage predominantly in frontoparietal regions (9/20 subjects), or stereotactic EEG (sEEG), predominantly in deep temporal lobe regions (amygdala, hippocampus, insula; 11/20 subjects). ECoG and sEEG activity was recorded, deidentified, and stored at the same time as behavioral data. Data were collected using Tucker-Davis Technologies, Nihon-Kohden, or Natus systems. Data processing was identical across all sites: channels were amplified x10000, analog filtered (0.01 - 1000 Hz) with > 1kHz digitization rate, re-referenced to a common average offline, high-pass filtered at 1.0 Hz with a symmetrical (phase true) finite impulse response (FIR) filter (∼35 dB/octave roll-off). Behavioral data were simultaneously collected using a PC laptop running Python (v.2.7) and PsychoPy (v.1.85.2) and synchronized with a timed visual stimulus (trial start) recorded by a photodiode through an analog input to the electrophysiological system.

### 2.3 Behavioral task

We probed risk-reward tradeoffs using a simple gambling task described previously (Saez et al., 2018). Briefly, subjects chose between a safe bet ($10, fixed) or a gamble for potential higher winnings (between $15 and $30). Gamble win probability varied per trial based on an integer between 0-10 shown at game presentation. Trials with 0 or 10 carried no risk as the outcome was certain and we used them as ‘catch’ trials to evaluate whether our model is decoding risk, rather than just choice selection independent of risk (Supplemental Data Figure 4). After choice (t = 550ms post-choice, Figure 1A), a second number (also 0 - 10) is revealed. The gamble results in a win if the second number is greater than the first, and ties were not allowed, therefore, a shown ‘2’ had a win probability of 80% and an ‘8’ had a win probability of 20%. Both numbers were randomly generated using a uniform distribution. Location of safe bet and gamble options (left/right) were randomized across trials. Subjects played 10 practice trials, repeated as many times as necessary, to ensure they had full knowledge of the (fair) structure of the task prior to game play (200 trials). Timing is summarized in Figure 1A. Trials started with a fixation cross (t = 0), followed by a game presentation screen (t = 750ms). Subjects had up to 8s to respond. Gamble outcome presentation appeared 550ms after button press (choice) on each trial regardless of choice. A new round started 1s after outcome reveal. The experimental task typically lasted 12-15min. This gambling task minimized other cognitive demands (working memory, learning, etc.) on our participants while allowing us to test for decision-making under risk. Behavioral performance was assessed by examining the proportion of trials in which the subject chose to gamble as a function of win probability; the proportion of risky trials was calculated for each win probability value (0-100% in 10% increments) and fit with a logistic curve. Subjects gambled more often as the win probability increased. As a control for behavioral data quality, we excluded subjects in which a logistic function did not appropriately fit the relationship between percentage of gambles and win probability (p<0.05, logistic fit). Results examining the correlation of differences in power between safe and gamble bets from these subjects playing the same task were published previously (Overton et al., 2025).

**Figure 1.**
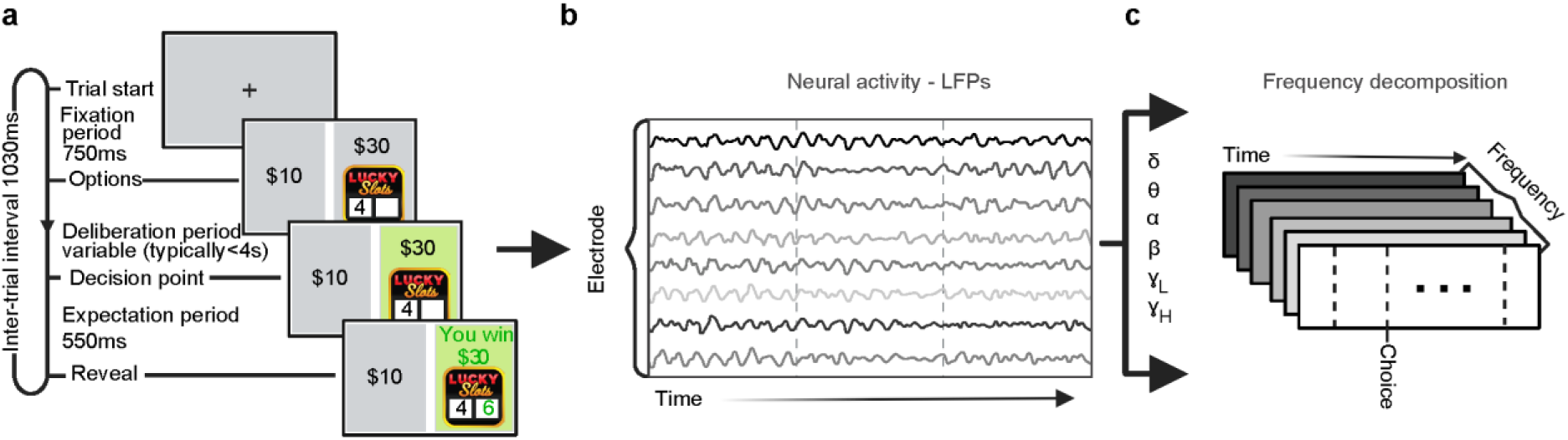
Task and Data Collection. **A.** Subjects played a gambling game in which they chose between a safe $10 reward and a risky gamble between a $30 reward or a $0 reward. Gamble win probability varied parametrically on a trial-by-trial basis (0-100%, 10% increments). **B.** Local field potentials (LFPs) from iEEG electrodes in several brain regions were cleaned to remove noisy channels and epochs, downsampled, filtered and referenced to a common average. (Overton et al., J Neurosci under revision). **C.** Data were then windowed around the time of choice for each trial and decomposed into power across frequency bands (delta [1-4Hz], theta [4-8Hz], alpha [8-12Hz], beta [12-30Hz], gamma [30-70Hz] and high-frequency activity [HFA; 70-200Hz]. One second is removed at the beginning and end of each trial segment, channels were z-scored over time to correct for the 1/f profile of neural activity.

### 2.4 Anatomical localization

Electrode localization was based strictly on clinical criteria for each subject, 9/20 had electrocorticography (ECoG) grids, predominantly in orbitofrontal, lateral prefrontal, and parietal regions, whereas 11/20 had stereotactic EEG (sEEG) coverage, predominantly of deep temporal lobe regions (amygdala, hippocampus) (Table 1). For each subject, a pre-operative anatomical MRI (T1) image and a post-implantation CT scan was collected. The CT scan allowed identification of individual electrodes but offered poor anatomical resolution, making it difficult to determine their anatomical location. Therefore, the CT scan was realigned to the pre-operative MRI scan following a previously described procedure^79^. Briefly, both the MRI and CT images were aligned to a common coordinate system and fused with each other using a rigid body transformation. Following CT-MR co-registration, we compensated for brain shift, an inward sinking and shrinking of brain tissue caused by the implantation surgery. A hull of the subject brain was generated using the FreeSurfer analysis suite, and each grid and strip was realigned independently onto the hull. This step was necessary to avoid localization errors of several millimeters common in ECoG subjects. Subsequently, each subject’s brain and the corresponding electrode locations were normalized to a template using a volume-based normalization technique and snapped to the cortical surface^79^. Finally, the electrode coordinates were cross-referenced with labeled anatomical atlases (Brainnetome atlas) to obtain the gross anatomical location of the electrodes, verified by visual confirmation of electrode location based on surgical notes. Electrodes across a broad set of regions known to be involved in reward-related behavior were used for analysis including lateral prefrontal cortex (LPFC; 391 electrodes from n=19 subjects), orbitofrontal cortex (OFC; 193, n=18), cingulate cortex (CC, 84, n=13), hippocampus (HC; 65, n=13), amygdala (Amy; 32, n=11), insula (Ins; 46, n=8), precentral gyrus (PrG; 108, n=11), postcentral gyrus (PoG; 88, n=9), and parietal cortex (PC; 78, n=8) and white matter (WM; 258, n=13). To avoid biasing the models based on whether the selected electrodes represented choice information, we included all electrodes in these analyses unless otherwise noted (Table 1).

**Table 1.** Count of electrodes per patient and region. Electrode counts for each subject broken down by region. The number of electrodes in each anatomical regions for each patient in our final sample of n=20 subjects, as well as the total count of electrodes per ROI across all patients and the number of patients with coverage for each ROI.

| Patient | LPFC | OFC | Cingulate | Amygdala | Hippocampus | Insula | Precentral Gyrus | Postcentral Gyrus | Parietal | Total electrode number |
| --- | --- | --- | --- | --- | --- | --- | --- | --- | --- | --- |
| p01 | 0 | 1 | 0 | 0 | 1 | 0 | 0 | 0 | 13 | 15 |
| p02 | 24 | 5 | 0 | 0 | 0 | 0 | 7 | 8 | 2 | 46 |
| p03 | 55 | 0 | 0 | 0 | 0 | 0 | 47 | 36 | 22 | 160 |
| p04 | 7 | 49 | 6 | 1 | 7 | 0 | 0 | 0 | 0 | 70 |
| p05 | 30 | 61 | 3 | 0 | 0 | 5 | 14 | 8 | 5 | 126 |
| p06 | 29 | 7 | 0 | 0 | 0 | 0 | 10 | 15 | 24 | 85 |
| p07 | 41 | 11 | 0 | 0 | 0 | 0 | 11 | 3 | 2 | 68 |
| p08 | 26 | 10 | 2 | 1 | 8 | 0 | 9 | 6 | 4 | 66 |
| p09 | 15 | 14 | 0 | 0 | 0 | 0 | 1 | 2 | 0 | 32 |
| p10 | 8 | 3 | 2 | 6 | 6 | 0 | 1 | 0 | 0 | 26 |
| p11 | 11 | 0 | 11 | 4 | 3 | 3 | 2 | 6 | 6 | 46 |
| p12 | 15 | 5 | 4 | 5 | 9 | 5 | 0 | 0 | 0 | 43 |
| p13 | 9 | 3 | 1 | 4 | 0 | 0 | 0 | 0 | 0 | 17 |
| p14 | 15 | 4 | 4 | 1 | 4 | 0 | 0 | 0 | 0 | 28 |
| p15 | 11 | 3 | 8 | 5 | 6 | 0 | 0 | 0 | 0 | 33 |
| p16 | 12 | 4 | 3 | 2 | 10 | 1 | 3 | 0 | 0 | 35 |
| p17 | 11 | 2 | 8 | 2 | 7 | 6 | 0 | 0 | 0 | 36 |
| p18 | 29 | 6 | 18 | 0 | 1 | 7 | 3 | 4 | 0 | 68 |
| p19 | 38 | 4 | 14 | 1 | 2 | 14 | 0 | 0 | 0 | 73 |
| p20 | 5 | 1 | 0 | 0 | 1 | 5 | 0 | 0 | 0 | 12 |
| **total per ROI** | **391** | **193** | **84** | **32** | **65** | **46** | **108** | **88** | **78** | **1085** |
| **# patients with ROI coverage** | **19** | **18** | **13** | **11** | **13** | **8** | **11** | **9** | **8** | **110** |

### 2.5 Electrophysiological analyses

#### 2.5.1 Quality control and preprocessing

As part of a study to regress change in power across frequency band with behavior (*Multi-Areal Neural Dynamics Encode Human Decision Making*, n.d.), LFPs were cleaned as follows. Epileptogenic channels and channels with excessive noise (low signal-to-noise ratio, 60 Hz line interference, electromagnetic equipment noise, amplifier saturation, poor contact with cortical surface) were identified and deleted. Data from 1317 out of 1452 electrodes were artifact-free and included in subsequent analyses. Each epoch of data was visually inspected to exclude epochs with aberrant or noisy activity (typically <1% of datapoints). Data analysis was done using custom scripts written in MATLAB and Fieldtrip toolbox^80^. Data for each channel were downsampled to 1KHz. Each channel was lowpass filtered (200Hz), highpass filtered (1Hz), and notch filtered (60Hz and harmonics) to remove line noise, and downsampled to 1 kHz if necessary. Electrode channels were re-referenced to a common average reference of all electrodes in each strip/grid. Even though bipolar derivations or white matter referencing are often used for sEEG electrodes, we opted to use a single (CAR) re-referencing strategy for both ECoG and sEEG electrodes for analytical consistency. Trials were epoched to the time of decision using a window starting 4s before choice and ending 3s after choice. Time-frequency representations (DPSS taper method) were plotted for each region and subject (averaged across electrodes and trials) and visually inspected for artifacts after the leading and trailing 1s of data were discarded to remove edge effects.

#### 2.5.2 Time-frequency representation of neural activity (bandpass estimates)

To assess the role of individual oscillatory bands, the LFP was decomponsed into canonical, discrete activity bands: (delta, δ [1-4Hz]; theta, θ [4-8Hz]; alpha, α [8-12Hz]; beta, β [12-30Hz]; low gamma, ɣ_L_ [30-70Hz]; hi gamma, ɣ_H_[70-200Hz]) for each electrode for each subject using the Filter-Hilbert method. Briefly, power in the 6 bands was calculated by applying a Butterworth bandpass filter (order 3 for delta and order 4 for all other power bands) and Hilbert transform and multiplying the resultant complex signal by its complex conjugate^81^. As before, one second is removed from the beginning and end of the data to reduce edge effects. Prior to dimensionality reduction for classification, power data for each trial and channel was z-scored over the time dimension within each band to correct for the 1/f profile of neural activity.

### 2.6 Decoding Models

Four different models were compared to assess how the frequency band activity encoding choice. Two different dimension reductions techniques: Principal component analysis (PCA) and linear dynamical systems (LDS) modeling were used. For each, two different classifiers were used: a simple Euclidean distance (ED) classifier that compared the single trial response to the average response of all other trials separated by class (gamble or safe bet) and dynamic time-warping (DTW) that generates a distance metric that is capable of accounting for trial-by-trial variation in time of neural activation through flexible stretching or shrinking of the time dimension. How these dimensional reduction techniques and classifiers were used is described next.

#### 2.6.1 Dimension Reduction

Principal components analysis (PCA) was used to assess the covariance structure of the data. First, we examined the redundancy of the data across electrodes within region for each frequency band. PCA was performed separately on each frequency band for all of the electrodes in a single region. The amount of variance carried by each PC was assessed (refer to Supplemental Data Figure 1). Second, to identify the optimal window of time and relevant frequencies of interest that encode choice, the variance captured when PCA was used to assess the covariance across all regions was examined using a scree plot and the optimal number of PCs was selected as the dimensionality of the system (Figure 2A). Then, the performance of the Euclidean distance classifier (ED, see below) on the first 200 ms before choice was calculated and this was repeated by increasing the size of the window by 200 ms increments up to 3 s prior to choice (Figure 2B). This was done separately for each subject and separately for each frequency band using leave-one-out cross-validation (see ED classifier, below). Lastly, the performance of PCA using the two different classifiers (PCA-ED and PCA-DTW) were compared (Student’s t-test).

**Figure 2.**
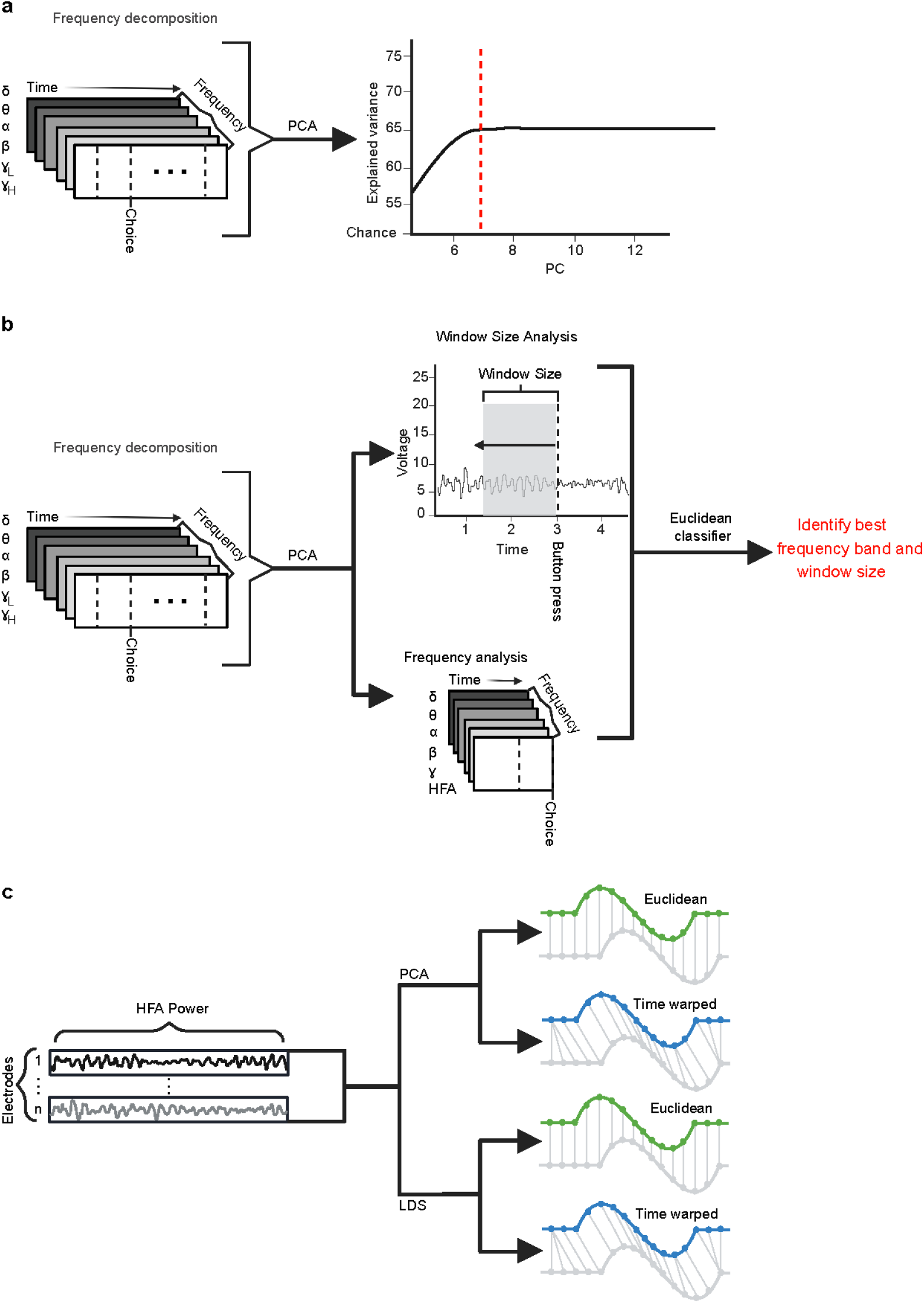
Feature engineering and model development. **A.** Principal components analysis was used to reduce the dimension of the data and identify features that conveyed the most information about choice selection. **B.** The first PC from each region for each frequency band was used to assess the optimal window size and most relevant frequency for decoding choice. **C.** The selected features were used to compare and contrast four linear models: PCA followed by a simple Euclidean distance (ED) classifier or a dynamic time warped (DTW) classifier and Linear Dynamical Systems model followed by the two classifiers.

In addition to PCA, linear dynamical systems (LDS) modeling was employed. LDS models allow characterization of the development of neural activity through time, and thus may be better suited for identifying how neural computations unfold during the task(Ghahramani & Hinton, 1996a, 1996b). LDS has been used to model the dynamics of neural activity (Sani et al., 2018, 2021), but mainly in the motor cortex (Disse et al., 2023; Kao et al., 2015, 2017; Orsborn et al., 2014; Shanechi et al., 2014; Shenoy et al., 2011, 2013a). LDS reduces the dimensionality of the data by learning a set of model parameters using expectation-maximization likelihood to identify a set of low dimension latent variables (LV) that capture the dynamics of the system such that the dynamics of the neural activity at the current time can be predicted from the dynamics at the prior time (Buesing et al., 2012). LDS models have proven to be effective in characterizing the behavior of neural networks and predicting the response of neurons to external stimuli . In particular, it allows us to leverage the high temporal resolution of iEEG recordings to better understand the neural system dynamics.

Similar to PCA analysis above, the appropriate dimensionality of the dynamical system model (number of LVs) was determined empirically by learning new model parameters for a range of dimensions (1-15) and examining the predictability of the neural dynamics. The dimension of the system was determined by the dimension that did not further improve the predictability of the neural dynamics. The dimensionality value ranged from 5 to 12 across subjects. Each dimension of the LDS model defines a latent variable that describes how the dynamics unfold and these LVs were exploited to further probe the underlying dynamics of choice processing.

#### 2.6.2 Classifiers

Two different classification techniques were employed to decode trial-by-trial subject choices from the neural data prior to button press (t=0): a simple Euclidean Distance (ED) classifier and a Dynamic Time Warping (DTW) approach. ED is computationally efficient and shown to be as effective as linear discriminate analysis (LDA) at classifying neural activity in response to different events (Foffani & Moxon, 2004). Briefly, we created separate neural trajectory templates for safe bets and gambles by averaging the reduced-dimensional neural features (see results) across trials, on a per subject basis, leaving out a single test trial. The test trial was left out from both the dimensionality reduction step and this averaging procedure. After the templates were calculated, the overall Euclidean distance (sum of the square differences for each bin (time) and each PC or LV dimension), between this trial and the average safe bet/gamble templates was calculated with that trial left out. The predicted choice was then assigned to the closest template in Euclidean space. If the predicted choice matched the true, behaviorally expressed choice (either safe bet or gamble), the trial was correctly classified. This procedure was repeated for all trials in our sample. Decoding accuracy, or performance, of the model was defined as the percentage of correctly classified trials, calculated on a subject-by-subject basis using balanced accuracy by calculating performance separately for each class (gamble and safe bet) and then averaging them.

While an efficient classification method, the ED classifier’s fixed template matching and temporal comparison may not be the best approach for decoding complex neural processes. We attempted to improve on the ED classifier by using dynamic time-warping (DTW), a distance metric that is capable of accounting for trial-by-trial variation in time of neural activation through flexible stretching or shrinking of the time dimension. Briefly, DTW seeks to minimize differences between two temporal sequences (in this case, the individual trial and each of the two templates) while allowing some flexibility in the matching. Specifically, instead of matching point-by-point, temporal warping that respects the sequence of points in each time series is allowed. DTW does this by using dynamic programming to calculate whether one or neither of the time series should be stretched in order to minimize the Euclidean distance between the two time series. We used the Matlab implementation of dynamic time warping (`dtw`). We hypothesized this additional temporal flexibility would result in classification improvement if the timing of neural activity supporting subject choices varies from trial to trial, but the underlying computations are the same. In brief, similar to the approach for Euclidean distance classifier, we leave out one trial and generate the template for safe and gamble trials by averaging over the trials. We then allow DTW to quantify the distance between the single and each of the templates separately, except here, we allow for time warping. The warped single trial is then assigned the class of the warped template that it is closest to. If the classification matches the participants choice on that trial then the trial is successfully classified. For DTW we used the same leave-one-out procedure with balanced accuracy as the ED classifier with the average for both safe and gamble trials warped separately with each single trial to identify the best match.

To identify bias in the neural data that may account for performance above the expected 50% chance performance, we randomly swapped the labels (gamble or safe bet) for each trial and assessed bootstrap performance. The distribution of performance provides an empirical assessment of chance performance for each subject. This is especially important here because in this task, subjects are more likely to select a gamble than a safe bet which could bias classifier performance. The bootstrap performance across subjects was almost exactly 50%, as expected (50.1±0.002%).

### 2.7 Contribution of brain areas/subcircuits to decoding accuracy

To examine the contribution of individual brain areas and subcircuits to decoding, the performance of the model (e.g. decoding accuracy) was calculated after progressively adding individual regions/subcircuits. We started by calculating performance using only the region with the worst individual classifier performance and iteratively added regions in order of increasing performance until all regions were included, separately for each subject. Because a single region might not have a significant impact on choice outcome, we also examined the contribution of subcircuits, by the same analysis using subcircuits (prefrontal: orbital frontal cortex, lateral prefrontal cortex and cingulate cortex), frontoparietal: precentral gyrus, post central gyrus and parietal cortex, and limbic: amygdala, hippocampus and insula) previously defined (Overton et al., 2025) instead of regions.

### 2.8 Latent variable trajectories and subspaces

We examined the latent variables (LVs) from the LDS model in order to probe the underlying dynamics of the neural activity. For demonstration purposes (refer to Figure 5), three of seven LVs from one subject that carried the most amount of information about choice were selected (refer to results). The selected LVs for gamble and safe bet trials were averaged separately and plotted against time and in 3D LV space. To confirm our observation that neural trajectories for safe and gamble trials converged to separate subspaces, we applied two approaches across all subjects.

First, the Lyapunov exponents for safe and gamble trials were estimated. The Lyapunov exponent measures the rate at which two similar, multi-dimensional time series diverge from each other as time progresses(Pesin, 1977) with positive values indicating that the trajectories are getting farther apart over time, e.g. more chaotic, and negative values indicating the trajectories are getting closer together over time, or converging, potentially towards an attractor. For each of the 7-dimensional trajectories, for each trial, we calculated the rate of divergence of the LV between trials within trial type to estimate the Lyapunov exponent over time separately for each subject. The temporal evolution of the exponents was averaged across subjects to evaluate whether, on average across subjects, the Lyapunov exponent declined towards negative values, indicating an attractor-like subspace.

Second, we examined how the trajectories for individual trials diverged from each other as the time of choice selection neared, separately for safe and gamble trial. The Matlab implementation of convex hull was used (`convhull`) to quantify the subspaces defined by the safe or gamble trajectories. At 100 ms intervals prior to choice, the LV data points from all of the safe or gamble trials separately were used to generate a safe hull and a gamble hull. Then the percentage of data points, separately for each trial, that were contained in both hulls was calculated to assess the overlap.

## 3. Results

Twenty medication refractory epilepsy subjects whose neural activity has previously been studied were included in this analysis (Overton et al., 2022; Saez et al., 2018). Local field potential data were recorded postoperatively from intracranial electroencephalographic (iEEG) electrodes while subjects played a gambling task^17^, during their stay at the epilepsy monitoring unit. For each trial, subjects could choose between a safe prize or a risky gamble with varying win probabilities (Saez et al., 2018). As expected, subjects gambled more often in trials with higher win probability, to maximize reward(Doll et al., 2012; Niv, 2009). LFPs from all electrodes were transformed into the magnitude of the power for 6 frequency bands (see Methods) and changes in power with time prior to choice were used to understand how these features of the data contributed to information about the subject’s choice. LFPs were recorded from several brain regions frequently targeted for clinical monitoring in patients, including orbitofrontal, lateral prefrontal, cingulate and parietal cortices, pre- and postcentral gyri, hippocampus and amygdala.

### 3.1 Covariance structure reveals role of high frequency activity for encoding choice

To assess the role of oscillatory activity, PCA was performed on the data from electrodes within a region separately for each power band, separately for each region. The first PC carried the majority of the variance (>50%) for all regions for each frequency band, suggesting a low dimensional system was a good representation (Supplemental Data Figure 1). In fact, more than 90% of the variance could be captured by less than 5 PCs (Supplemental Data Figure 1) suggesting that, within region, most of the variance is linear. Upon visual examination, the first PC was a good representation of the average power in each region for each frequency band.

To assess the dimensionality of the system across regions, PCA was repeated using all the electrodes separately for each frequency bands. More than 80% of the variance was explained by 7 PCs across all frequency bands (Figure 3A). To identify which frequency bands conveyed the most information about choice, a simple Euclidean distance classifier (ED) was used to access when the information was represented (Figure 3B), using a leave-one-out validation strategy and balanced accuracy. As expected (Saez et al., 2018), both low and high gamma activity separately conveyed significant amounts of information (>55% performance) with data from all of the other frequency bands only slightly better than chance (bootstrapped random performance 50.1±0.002%). Interestingly, the low gamma band activity peaked immediately prior to choice and increasing the window actually disrupted this information. The high gamma activity, on the other hand, contained the most information about choice, peaking when a 1 s window was used, suggesting that information about choice accumulated during the trial since the average duration was greater than 1.5s (average reaction time 1.52 ± average std dev 5.88s). Thus, in subsequent analyses using PCA, both low and high gamma band activity were included, reducing the number of features for each subject from the varying number of electrodes per region by six frequency bands to 7 PCs.

**Figure 3.**
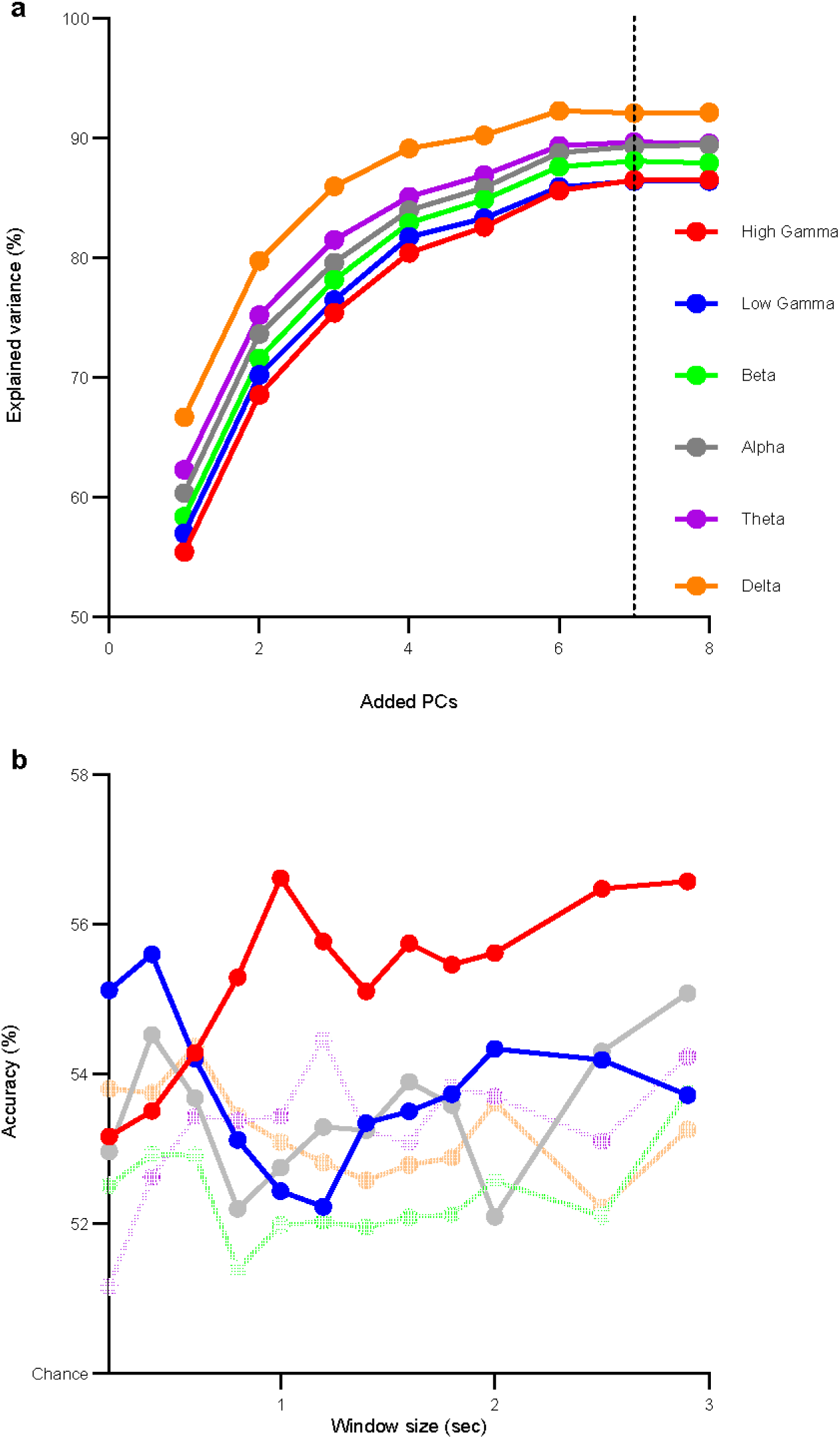
Covariance structure of power changes prior to choice reveals neural features that encode choice. **A.** The variance within each region for each frequency band is captured by the first 7 principal components, as denoted by the dotted black line. **B.** HFA activity including activity in both low (30-70Hz) and high (>70 Hz) gamma power band, provides most of the information about the subject’s choice in the second before choice selection.

**Figure 4.**
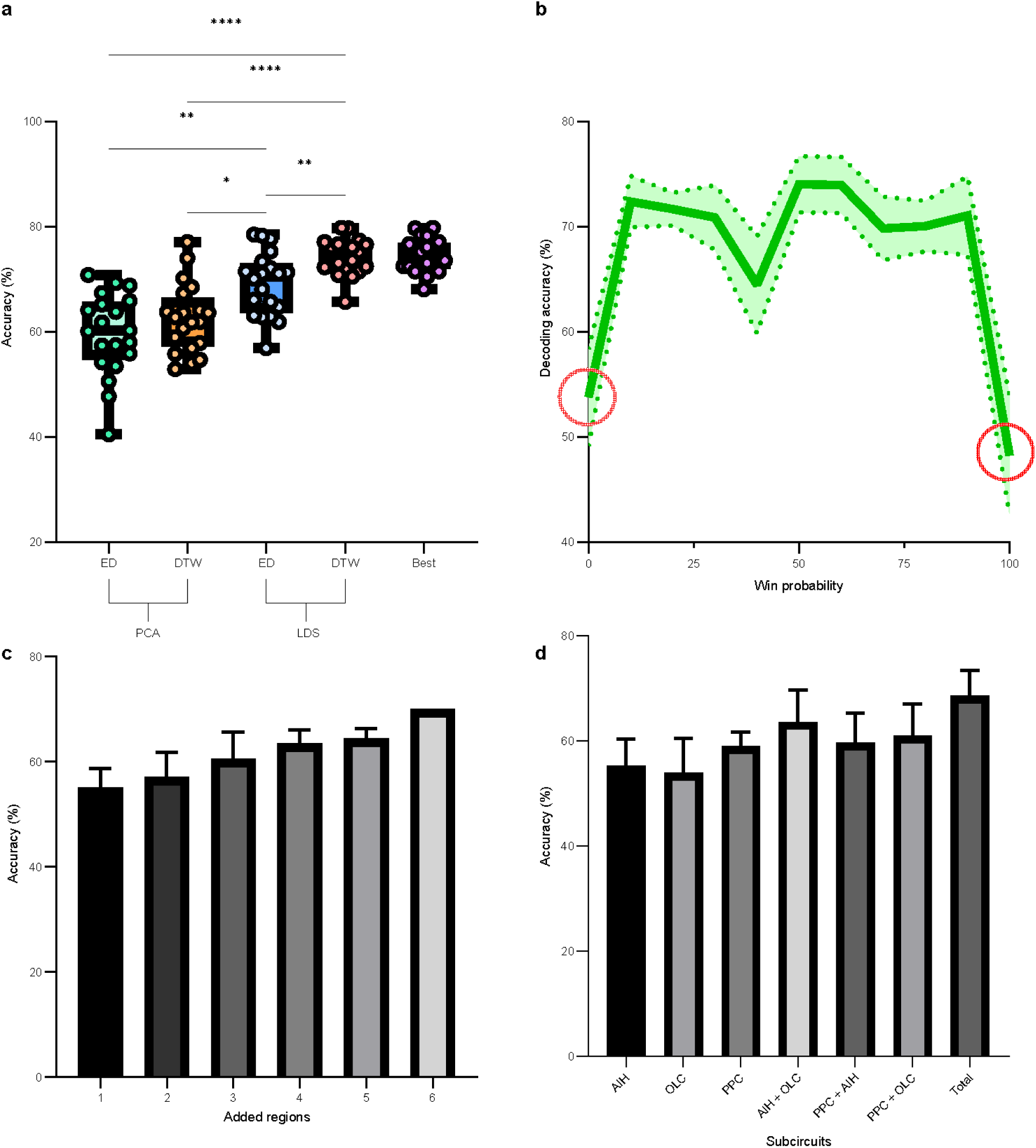
Linear dynamic system modeling. **A.** The LDS-based models outperformed the PCA-based models regardless of whether the simple Euclidean Distance classifier (ED) or the dynamic time warping (DTW) classifier was used. When the best performance for each subject was selected from each of the model, there was no difference between LDS-DTW and best performance suggesting that LDS-DTW will work as the best model for most subjects. **B.** Model performance fell to chance when no real option was offered (i.e. win probability equal to 0 or 100, denoted by red circles). However, when a choice is offered, model performance is the same regardless of the win probability. **C.** Accuracy of the model increases linearly as the number of regions included increase. **D.** Further, dropping any subcircuit from the model does not impact accuracy. Comparisons were made using one-way ANOVA with Tukey post-hoc when appropriate. *p < 0.05, ** p < .01, ****p <.0001.

**Figure 5.**
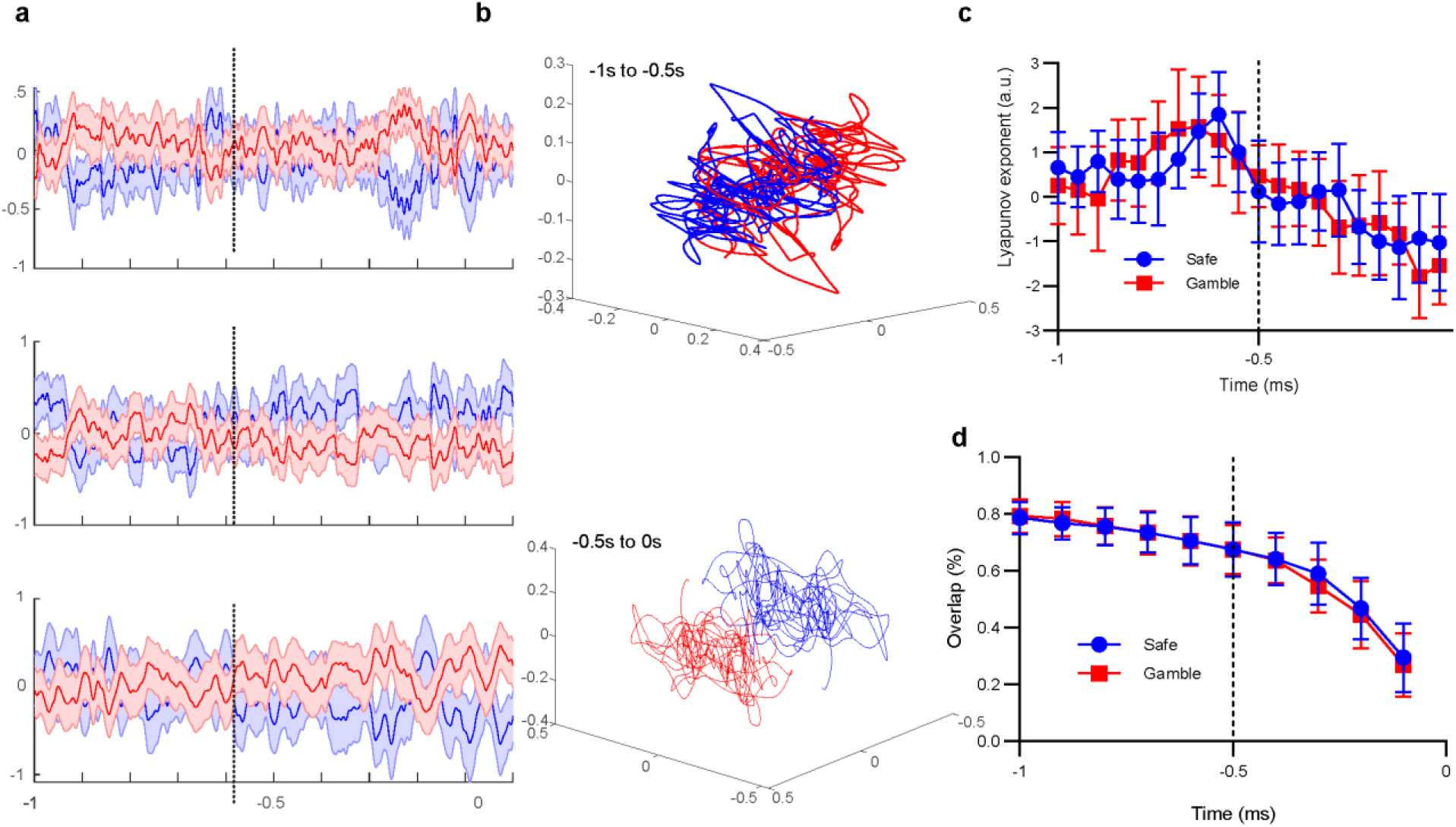
Linear dynamic systems modeling provides insight into decision-making behavior. **A.** Latent variables (LV) plotted as function of time before choice. The 3 LVs plotted here have the best individual classification performance of the 7 possible LVs per subject and show increased separate between gamble and safe choices at approximately 500 ms prior to choice as denoted by the dotted black line. **B.** The same 3’ LVs plotted for one subject with the best overall performance. Top plot shows the time between 1 s and 0.5 s prior to choice. Bottom shows the 0.5 s before choice. **C.** For both safe (top) and gamble (bottom) trials, the Lyapunov exponent is positive early during deliberation but then becomes negative approximately 0.5 s before button press (time=0) denoted by dotted black line. **D.** The overlap between convex hulls over the latent variables of the safe trials and gamble trials generated at 100 ms intervals prior to choice decreases as the time of choice approaches and transitions from a linear change to a supra-linear change at 500 ms prior to choice denoted by dotted black line.

This simple ED classifier, using 7 PCs derived from high frequency activity (HFA: both high and low gamma) had an average performance of 59.6±7.6% with performance ranging from just above chance (40.6%) to just over 70% (70.8%). Given the variability in the duration of each trial, we suspected that while the underlying covariance structure might be the same, the neural computations performed during short duration trials would occur more quickly while for longer duration trials, the computation would evolve more slowly. To test this, we compared this ED classifier with one that utilized dynamic time warping (see Methods). DTW had a modest yet significant improvement in performance (62.5±6.36%) over the strict average response of the ED classifier (p<0.05, Student’s t-test one-sided, paired), suggesting some trial-to-trial variability in the speed at which the computation was performed.

### 3.2 Neural population dynamics exhibit a low-dimensional structure that discriminates choice

LDS characterizes ongoing neural dynamics by identifying the underlying latent variables (LVs) that describe how the dynamics at the current moment in time produce the dynamics at the next moment in time (see Methods). These LVs capture the linear component of these neural dynamics. We constructed these 7 LVs (see Methods) for each trial in the 1 s before choice and repeated the ED and DTW classification, again using a leave-one-out validation strategy and balanced accuracy.

LDS outperformed PCA regardless of the classifier (ANOVA F(3,3734)=4.17; p < 0.005; Figure 4A). The optimal decoding strategy, LDS followed by DTW, accurately classified choice 74.3±3.4% across all subjects (Table 2) and its performance was close to the maximal or best performance for all subjects (Figure. 4A “Best”, 74.6%±3.2%). Therefore, a low dimensional, linear dynamical model followed by a classifier that accounted for variation in temporal dynamics across trials was the best strategy for choice decoding.

**Table 2.** Decoder performance per patient. Performance (% accurately classified trials) for each model for each patient. Trials with no uncertainty (i.e., gamble win probabilities of 0% and 100%) are excluded. Data Table corresponds to Figure 4a. Abbreviations: linear dynamical systems (LDS), Principal Components analysis (PCA), Euclidean distance classifier (ED), dynamic time warping classifier (DTW). Best is simply the best performance of the 4 models.

| Patient | LDS+DTW | LDS + ED | PCA+DTW | PCA + ED | Best |
| --- | --- | --- | --- | --- | --- |
| p01 | 65.7% | 68.1% | 63.8% | 60.3% | 68.1% |
| p02 | 76.5% | 61.9% | 57.9% | 67.4% | 76.5% |
| p03 | 72.5% | 62.9% | 61.9% | 70.8% | 72.5% |
| p04 | 76.1% | 56.9% | 63.6% | 57.5% | 76.1% |
| p05 | 77.1% | 62.8% | 74.0% | 69.3% | 77.1% |
| p06 | 79.7% | 77.7% | 77.1% | 68.8% | 79.7% |
| p07 | 76.7% | 64.8% | 54.0% | 55.9% | 76.7% |
| p08 | 71.9% | 65.7% | 67.3% | 61.8% | 71.9% |
| p09 | 77.4% | 78.5% | 68.5% | 58.0% | 78.5% |
| p10 | 79.8% | 71.3% | 60.7% | 53.5% | 79.8% |
| p11 | 71.5% | 66.1% | 62.8% | 65.9% | 71.5% |
| p12 | 75.2% | 73.4% | 56.9% | 64.1% | 75.2% |
| p13 | 73.6% | 72.1% | 59.0% | 50.6% | 73.6% |
| p14 | 70.5% | 63.1% | 61.8% | 57.4% | 70.5% |
| p15 | 76.6% | 78.3% | 54.7% | 54.2% | 78.3% |
| p16 | 72.3% | 71.1% | 64.1% | 65.2% | 72.3% |
| p17 | 76.1% | 75.3% | 55.8% | 40.5% | 76.1% |
| p18 | 72.4% | 70.1% | 63.6% | 60.1% | 72.4% |
| p19 | 73.1% | 71.1% | 70.2% | 63.8% | 73.1% |
| p20 | 70.8% | 71.5% | 52.9% | 47.7% | 71.5% |
| Mean | 74.3% | 69.1% | 62.5% | 59.6% | 74.6% |
| SD | 3.36% | 5.86% | 6.36% | 7.60% | 3.19% |

Importantly, the LDS+DTW model achieved above-chance decoding in all subjects, with a minimum decoding accuracy of 65.7%, indicating that it was robust to the variation in number of electrodes and anatomical localization across subjects (refer to Table 1). Decoding accuracy was as high as 79.7% for one subject, p06 (Table 2). There were no differences between decoding performance between safe and gamble trials (paired t-test, t(19)=0.6152, p > .05). Importantly, model performance depended on the win probability of the gamble trials, with the model performance falling to chance levels when there was no risk associated with the choice (i.e. win probability either 0% or 100%). In contrast, model performance was similar for all risky choices (win probability between 10% and 90%; Figure 4B. Furthermore, even when the choice was straight-forward (win probabilities of 10%, 20%, 80% or 90%), information about the subject’s choice selection was represented by the neural data as well as the choice on more difficult trials (30%-70% win probability). These differences in decoding accuracy could not be accounted for by differences in reaction time or gamma power (t-test, Bonferroni corrected p>0.025) between trials with no risk and trials with varying degrees of risk. Therefore, we excluded trials with no uncertainty in the outcome (gamble win probability = 0% or 100%) from model results.

These results provided confidence in our model to further assess the relative contribution of individual brain areas to classifying choice. Model training and classification was performed as individual regions were iteratively added to the decoding model starting with the least informative region (Figure 4C). For individual regions, the marginal increase for each region added suggests that within subject, no particular region conveyed a majority of the information (ANOVA F(1.889, 35.89)=25.44). To account for the possibility that known subcircuits that these regions are a part of ((Overton et al., 2025) and see methods) convey significant information that is not observable by looking at each region separately, we examined the information conveyed by subcircuits. Similar to the single region analysis, performance was not dependent on whether any specific subcircuit was included in the model (Figure 4D). These results further our understanding of the distributed nature of the encoding of choice demonstrating that although specific brain regions may contain specific information about choice-related variables including value ((Rich & Wallis, 2016)) win-probability ((Hampton et al., 2007)) and risk ((Schultz et al., 2008)), information about the final choice selection is shared among broad regions of the brain.

### 3.3 Latent variables reveal sub-second neural dynamics underlying choice

To gain further insight into the neural computations during deliberation prior to choice selection, the LVs for each trial were averaged separately for gamble and safe bet trials. Because some LVs may not reflect choice-related activity, for visualization purposes, we selected the three LVs whose representations during gamble trials were most different from safe bet trials, as assessed by their Euclidean distance, separately for each subject. Each of these LVs capture different components of the neural dynamics underlying choices. They separated repeatedly during the deliberation phase, albeit with different temporal dynamics, especially within 500ms of choice (Figure 5A and Supplemental Figure 2). To gain an understanding of these temporal dynamics during deliberation and their relationship to behavior, we examined the 3-dimensional LV projections for safe bet and risky gamble choice trials as they unfolded in time within the state space (Figure 5B and Supplemental Figure 3). First, we observed that the neural trajectories occupy a limited portion of the state space that captures all the data points prior to choice. Additionally, early in the deliberation period (−1 s to 500 ms prior to choice), safe bet and gamble trajectories overlapped as the single trial trajectories moved back and forth from one end of the state space to the other (Figure 5B top). As the decision point for choice approached (500 to 0ms pre-choice), safe bet and gamble LV trajectories diverged into non-overlapping subspace, suggesting a bistable system and the possible existence of separate attractors for gamble and safe bets (Figure 5B bottom and Supplemental Data Movie 1).

To confirm our observation that neural trajectories for safe and gamble trials have similar dynamics and converged to separate subspaces, we applied two approaches across all subjects (see Methods section 2.8). First, to assess the generalizability of this effect, we estimated and compared Lyapunov exponents separately for safe and gamble bets using all 7 LVs for each subject. Lyapunov exponents represent whether a system is chaotic or converging to some stable or dissipative subsystem over time. Specifically, Lyapunov exponents measure the rate at which two initially similar, multi-dimensional trajectories diverge from each other as time progresses^35^, with positive values indicating that the trajectories are getting farther apart over time and negative values indicating they are converging. For both gamble and safe bets trials, the Lyapunov exponents were calculated to measure the temporal rate of divergence separately for each subject and the result was averaged to examine temporal dynamics across subjects. On average, we observed that the value of the exponents early before choice (−1.0 s to - 0.5 s) was different from immediately before choice (−0.5 to 0.0 s; t(38) = 7.197, p < .0001). Moreover, on average, the exponents for both safe and gamble trials transitioned from positive to negative by 300 ms prior to choice, suggesting that the trajectories, across subjects, did in fact converge towards subspaces for gamble and safe bets as subjects neared a decision (Figure 5C). Thus, single trial trajectories first repeatedly visit the subspace for gamble and then safe bet, suggestive of deliberation. Then, the trajectory moves towards and spends more time in one or the other subspace, providing a network level view of how the brain can keep track of the accumulation of evidence for one or the other choice.

Second, to examine how these subspaces capture the neural dynamics for safe versus gamble trials evolve over time, we built convex hulls around the data points for safe and gamble bets separately at 100 ms intervals prior to choice and assessed the percentage of data points that were contained in both hulls (% overlap, Figure 5D). Consistent with the Lyapunov assessment, as we approached choice from 1 s prior, the percentage of data points with overlap decreased. The reduction in overlap accelerated as choice was approached (log fit r^2^ = 0.98) suggesting a rapid transition to the separate subspaces as the decision was made. For both approaches (Lyapunov exponent and convex hull) the transition to separate subspace occurs after 500 ms prior to choice, supporting our 3D visualization (see Supplemental Data Figure 2). Taken together, this network level view shows single trial trajectories repeatedly transversing subspaces to encode each choice followed by progressive separation to the subspace representing the preferred choice until sufficient information is accumulated for the participant to make a selection.

## 4. Discussion

Economic-based decision-making depends on coordinated activity across multiple brain areas, but how this distributed neural activity encodes information about choice in the human is not well understood. Our results provide several novel insights into the neural computations underlying human decision-making. First, for each frequency band, over half of the covariance of the power modulations can be represented by a single dimension supporting recent work that linear models capture a significant amount of neural activity (Gallego et al., 2020; Nozari et al., 2020, 2024). Second, the activity in HFA carries most of the information, which is consistent with prior studies showing that HFA is a good approximation of the underlying single neuron activity (Buzsáki et al., 2012; Ray et al., 2008; Ray & Maunsell, 2011) which often encode information about choice (Anders et al., 2017; Azab & Hayden, 2017, 2018; Blanchard & Hayden, 2014; Enel et al., 2020; Hayden et al., 2011; Pearson et al., 2009; Wallis, 2012, 2018; Wallis & Miller, 2003). Importantly, our decoding model fails when there is no risk associated with the gamble option (win probability of 0 or 100) but does similarly well in uncertain trials regardless of the win probability (10-90%). Therefore, the computations made during risky decisions are substantively different from those in non-risky decisions, potentially reflecting the fact that certain win/loss trials can be conceptualized as single option trials rather than value comparison trials.

Our results also show that allowing warping of the time prior to choice improves classification performance. This suggests that the shape trial-by-trial neural trajectories that are derived from the latent variables are similar, regardless of the duration of the trial. In addition, results show that LDS, which models the neural dynamics, outperforms PCA, which models the covariance. The neural dynamic of HFA, as noted above, reflect the underlying neuronal activity. These latent variables thus capture an emergent property of the neural activity that is shared across brain regions. Current recording technologies and analytical techniques have allowed for a deeper comprehension of the computations that populations of cortical neurons employ together as an ensemble (Gallego et al., 2017), during motor planning and execution (Churchland et al., 2007, 2010a; Michaels et al., 2016; Shenoy et al., 2013b), as well as during locomotion (DiGiovanna et al., 2016; Disse et al., 2023; Melbaum et al., 2022; Mimica et al., 2018) (Disse et al., 2023). A significant result of this work has suggested that characterizing the dynamical response of a neural population, rather than simply assessing covariance between neurons, can allow for better predictions of neural activity and, importantly, provide insight into the computations the brain performs during motor tasks. Therefore, to understand the computations undertaken by the cortex during postural control, it is necessary to characterize population dynamics.

Finally, these results provide insight into the likely importance of non-linear information in the data. The fact that the LVs from LDS converge to their own unique subspaces prior to choice, is highly suggestive of nonlinear activity since a single linear system cannot model two attractors. Moreover, the fact that choice for trials with win probabilities of 0 or 1 could not be decoded also supports that some of the information is likely encoded in a nonlinear representation of the data. Therefore, while noted above, a significant amount of information is carried by the linear model, nonlinear models are likely to capture more, and will, therefore, likely have better single trial decoding performance, as has been note for motor system (Abbaspourazad et al., 2024; Chapin et al., 1999; Christen et al., 2004; Song & Berger, 2010; Yu et al., 2008). Together, these data provide important insights into the computations the brain performs during decision-making.

Our approach relied strictly on linear dimension reduction techniques and linear classifiers. To date, almost all studies of the neural underpinnings of cognitive process use linear correlations and linear statistics to assess average differences in experimental classes, and they have provided important insights. Linear classifiers have been used in the past to understand neural computations associated with decision-making, including linear discriminant analysis (Avvaru et al., 2021; Basu et al., 2021; Provenza et al., 2019; Rich & Wallis, 2016; Rossi-Pool et al., 2017; Wallis, 2018). This is an initial approximation, since neural computations have a non-linear component. Nonetheless, linear approaches have three important advantages. First their impact on the data is easily interpretable and can be more easily tied back to the underlying neural activity. The fact that the linear component of the information was able to correctly classify a high proportion of single trials correctly suggests the models captured a significant amount of information about choice. Second, some of our principal results (low frequency activity across a single brain region can be captured by a one-dimensional model or high frequency activity carries most of the information) are strongly supported by earlier work in animal models, lending further confidence in the value of the model. Finally, our long-term goal is to develop closed-loop neuromodulation systems to aid those with pathological decision-making disorders (e.g. depression or addiction). A linear model can guarantee robustness and safety (Tóth, 2010). Nonetheless, more work needs to be done to understand what information is missing, preventing better performance, and non-linear models will likely shed insight into this in future studies. Combined with the ability to directly and precisely modulate brain activity using direct electrical stimulation^71–78^, these results open the door to the development of cognitive prostheses to modulate abstract brain states in the human brain to reduce the likelihood of pathological brain states (Saez & Gu, 2023).

Despite the fact that we restricted our analysis to low dimensional, linear models, their performance at correctly classifying the subject’s choice, or decoding accuracy, was high, especially considering our model produces trial-by-trial decoding. Therefore, regardless of the brain’s non-linear, higher order computations, substantial information is stored in a low dimensional linear manifold. The model performed significantly above chance in all subjects in our sample, despite differences in electrode location. Our decoding accuracy was significantly higher than comparable non-invasive decoding approaches. For example, Vickery et al (Vickery et al., 2011) used MVPA whole-brain approaches to decode both outcome (win/loss) information and choice (stay/shift) from distributed brain activity, but only achieved a mean accuracy of 52.6% for binary choices. A similar study using voxel-based techniques, found decoding accuracies as high as 64% in ACC(Hampton et al., 2007). A recent (2018) review reported that typical fMRI MVPA-based decoding accuracies for reward-related signals are a few percent points above chance(Kahnt, 2018). Our robust decoding performance is consistent with previous studies showing successful decoding of mental states from iEEG data(Kirkby et al., 2018; Sani et al., 2018; Thiery et al., 2020) and suggests that iEEG data provides a more sensitive measure of the information encoded about choice than non-invasive approaches. Therefore, despite the need to learn the specific parameters of the model for each subject, consistent encoding of choice information can be achieved by a low dimensional, linear model and supports their development to gain insight into human behavior.

### 4.1 Latent variables reveal neural computations underlying choice

Four results combine to provide insight into the neural computations underlying choice in this task. First, the model could not decode deterministic ‘catch’ trials (win probability equal to 0% or 100%) where the participant still needed to make a selection, but the risk was zero and, therefore, not a factor for choice selection. This suggests that the model identified neural computations that evaluated the risk underlying the choice selection. In fact, prior studies have shown significantly different behavioral responses to decision making when risk is present (Potenza et al., 2019; Schiebener et al, 2015). Moreover, certain brain regions, such as the parietal cortex, dorsolateral prefrontal cortex and anterior insula show changes in activation depending on whether a choice involves risk or not (Mohr et al., 2010; Wu et al., 2021). Our results clarify that not only are there behavioral differences but these changes in activation reflect fundamental differences in the neural computations when risk is involved with a decision.

Second, in contrast, performance across a broad range of win probabilities (10-90%) was similar, suggesting that the neural computations for easier choices are similar to those for more difficult choices. These likely involve calculating and comparing the expected utility of both options (safe bet and gamble) to guide choices. Third, the time warping algorithm (DTW) improved decoding performance, suggesting that decision-encoding had consistent yet time-varying trial-by-trial temporal dynamics that impact the ability of models to decode choice. Finally, a small number of latent variables captured broadly distributed, high dimensional neural activity, suggesting significant redundancy in the data. In fact, despite differences in anatomical coverage and surgical strategy from subject to subject, these models robustly decoded single trial activity to predict choice in every participant in the dataset. This redundancy is likely a necessary function of the system to bring together information from different types of calculations performed in different brain regions (e.g. expected value in OFC (Rich & Wallis, 2017)) in order for the subject to make the final selection, supporting prior studies that that specific information about regional computations is shared broadly across the brain (Overton et al., 2025).

Together, these results support two important hypotheses underlying choice selection in this type of task. First, while it is understood that within OFC, for example, neural activity scales with risk, the neural computations underlying risky choices as identified by the latent variables, are qualitatively similar, regardless of the amount of risk. A subject may be able to perform a low-risk trial (very high or very low win probability trials) faster, but the neural computations are the same and, importantly, provide the same information as more difficult trials with moderate win probabilities. Second, the neural information about risk that drives the final choice in this type of task is shared widely across the brain despite the fact that individual brain regions perform different calculations to assess different components of that overall decision (e.g. OFC = value). To confirm these hypotheses requires additional study that will likely include more complex tasks and non-linear decoding models. Nonetheless, the work presented here clarifies common computations underlying risky decision-making.

### 4.2 Attractor-like dynamics of power modulations underly deliberation

In latent space, the movement of the neural trajectories across the entire state space during deliberation suggests that the subject alternates between considering one choice and then the other.

This intuitively makes sense and is also consistent with linear discriminant analysis approaches to decoding choice during deliberation in animal studies (Rich & Wallis, 2017), demonstrating an important role for sub-second dynamics underlying choices that are suggestive of a dissipative system. This is the first view of such latent activity in humans. Importantly, both the changes in the Lyapunov exponent and the changes in overlap between the subspaces during deliberation were the same, suggesting that despite the fact that they converge to different subspaces, the time horizon of this convergence is similar for safe and gamble bets. Moreover, the fact that the neural trajectories were relatively chaotic more than 300 ms before choice but then accelerated to separate subspaces as choice selection neared suggests that once enough information to make the choice was collected, neural activity moves to a smaller space reminiscent of the coalescing of neural activity identified prior to movement in a delayed go-cue task (Churchland et al., 2010b). In that study, the data were recorded from pre-motor cortex and the final subspace was the same regardless of the choice the animal ultimately made. Here, we recorded across broad regions of the brain, including those that are part of a prefrontal (OFC/LPFC/CC), limbic (Amy/HC/Ins) and frontoparietal (PrG/PoG/PC) subcircuits and identified different subspaces for each option.

Further, the latent variables that best discriminated between gamble and safe bets suggest a view of deliberation wherein the neural representations underlying each option define separate attractors for that option, or a bistable system in the case of two options. During deliberation, this low dimensional representation of the neural dynamics traversed the state space multiple times. In fact, rapid transitions between the subspaces were observed for multiple trajectories separately, albeit with different dynamics, demonstrating that both safe bet and gamble subspaces were visited multiple times before a choice was made. This is similar to activity patterns observed in multi-electrode OFC recordings during deliberation in non-human primates that reflect the fast alternative evaluation of binary choices as in our task (Peixoto et al., 2021; Rich & Wallis, 2016).

Yet, as choice selection approached, the neural trajectories progressively converged to one of the subspaces. Importantly, this progressive separation of the neural dynamics required a higher dimensional space (multiple LVs) to observe. Moreover, this progressive nature suggests that the neural computations underlying deliberation evolve until a decision-threshold is reached, similar to non-human primate studies of evidence accumulation(Ditterich, 2006; Mazurek et al., 2003). Therefore, our results support work from animal studies about accumulation of knowledge on the one hand and consideration of options on the other and clarifies that during the deliberative process both rapid transitions considering each option and accumulation of knowledge about the ultimate choice are occurring simultaneously. Further, these latent dynamics suggest that these two processes can be represented by the same neural dynamics. This suggests that the neural computations underlying deliberation related to considering options and knowledge accumulation have similar underlying structure, but additional work would be needed to draw a firm conclusion. However, more work needs to be done to assign a causal relationship between these network level phenomena and behavior. Rather, we consider this state-space representation and the behavior of these neural trajectories as a view of the system that provides insight into how brain-wide networks gather information from populations of single neurons performing the specific computation. A majority of studies examining reward-based choice patterns use binary choices; our results open the door to the study of multi-option processes, since they predict that multiple subspaces would appear in the case of multi-option choices.

In summary, invasive iEEG recordings with high anatomical specificity, temporal resolution, and neurobiological detail, combined with economic probes of decision-making, and machine learning approaches provide insight into neural computations underlying decision-making behavior. Specifically, (1) information about decision-making is broadly distributed and carried by low dimensional, linear latent variables comprised of high frequency activity (>30Hz), (2) computations underlying easy versus more difficult choices are similar, as long as some uncertainty exists, (3) during deliberation these LVs move through large regions of the manifold, repeatedly visiting smaller subspaces that represent the choice options, and (4) the ability of the neural trajectories to separate into distinct subspaces prior to choice improved behavioral outcome and likely reflects the progression towards a decision threshold. Therefore, the models presented here suggest computations underlying single trial behavior involves rapid switching between options and knowledge accumulation toward a final decision are represented by similar neural dynamics.

## Acknowledgements

We would like to thank L. Nuñez, C. Meikle, and C. Foreman for help with data collection. We would especially like to thank the subjects for their willingness to participate in this research. We would like to acknowledge © Zoubin Ghahramani, University of Toronto for his code Version 1.0 01-Feb-98 (revised Mar-02) found here https://mlg.eng.cam.ac.uk/zoubin/software.html which we use in our code that can be found here: https://github.com/NeuralStorm/NeuralDynamicsEncodeChoice

## Contribution statement

IS, MH and RTK designed the research task; IS and JAO collected data; IS, JAO, LMP, AR, MPS, and KM analyzed data; RTK, EFC, JJL provided clinical supervision; IS, KAM wrote the paper.

