## Supplemental Material for "Neural dynamics encoding risky choices during deliberation reveal separate choice subspaces"

**Author list:** Logan M. Peters^1^, Alec Roadarmel^1^, Jacqueline A. Overton^3,4^, Matthew P. Stickle^1^, , Jack J. Lin^5^, Edward F. Chang^6^, Robert T. Knight^7,8^, Ming Hsu^7,9^, Ignacio Saez^10,11 ƚ^. Karen Anne Moxon^1,2 ƚ^ ^*^

To make clear how the principal components capture the variance, PCA was performed separately within each region for each frequency band. For each region and each frequency band, most of the explained variance was captured by a few components.

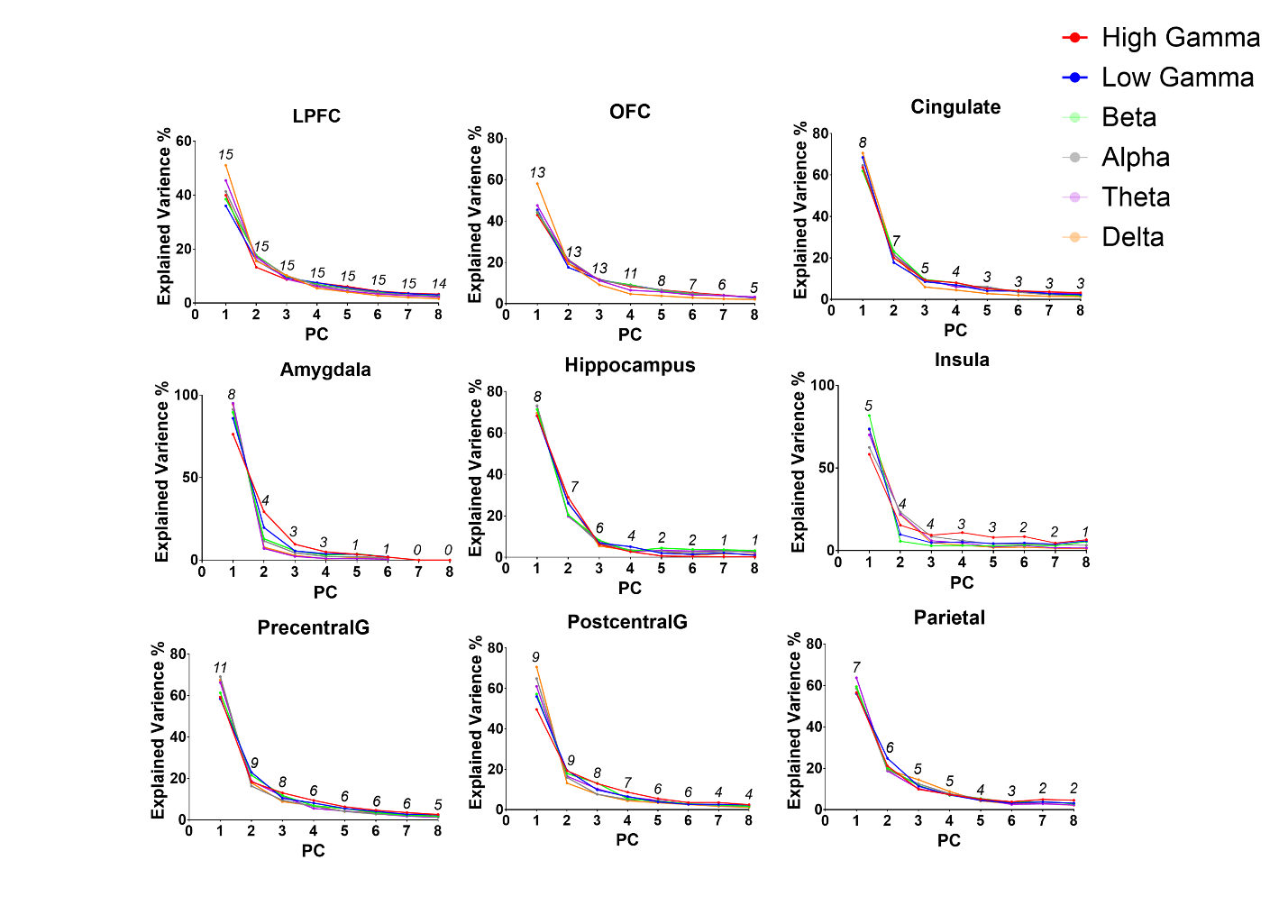

**Supplemental Data Figure 1. Covariance structure of power changes prior to choice reveals neural features that encode choice.** When principal components were constructed separatepy fro each region, more than 90% of the variance within each region for each frequency band is captured by less than 5 principal components.

To provide clarity regarding the variance in the latent variables across subjects, the three latent variables that best classify choice are plotted as a function of time (Supplemental Figure 2) and in 3-space (Supplemental Figure 3). Some subjects clearly show better separate between safe and gamble trials than others. This difference is manifest in the variability of the performance across subjects. Note that performance of the LDS models using the two classifiers (ED or DTW) was done using all 7 latent variables so these plots do not give a complete picture.

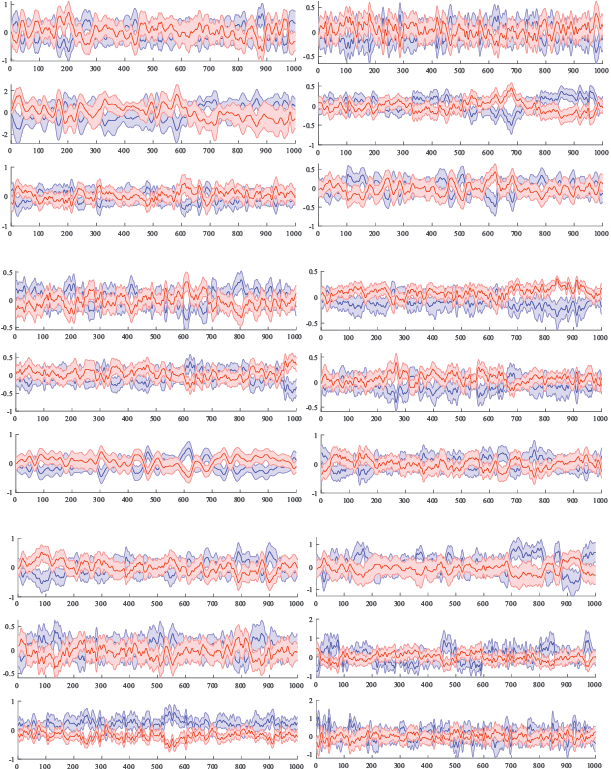

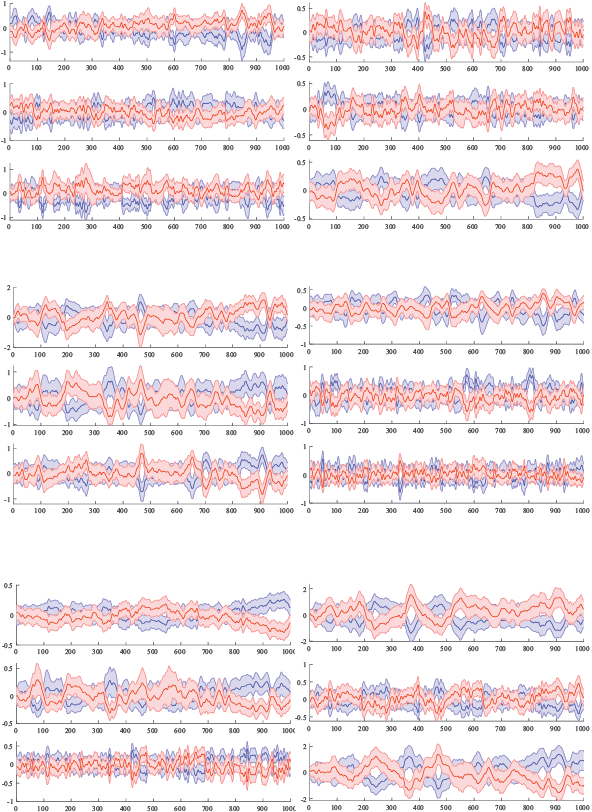

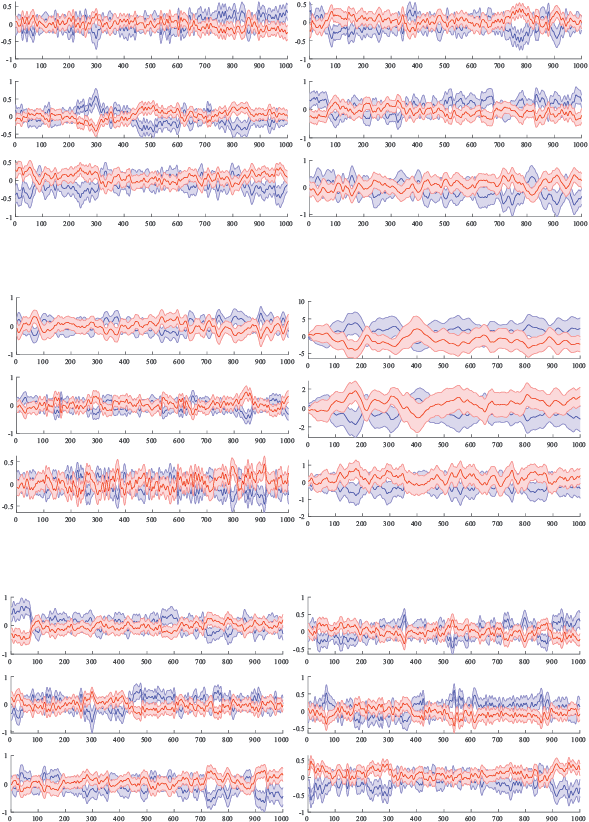

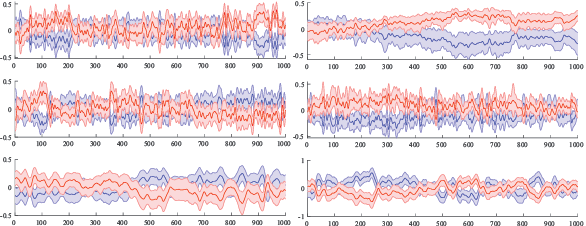

**Supplemental Data Figure 2.** The three latent variables that best separate gamble trials from safe trials plotted separately before choice for each subject. The latent variables for all gamble trials and all safe trials are separately averaged and graphed during the 1s before choice with shaded standard error. Red are gamble trials and blue are safe trials.

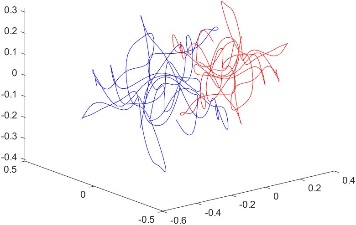

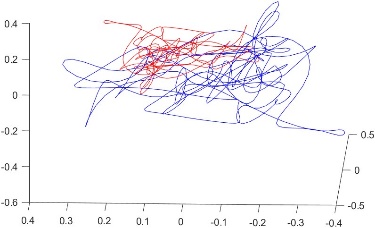

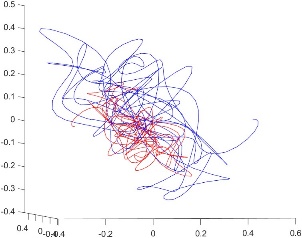

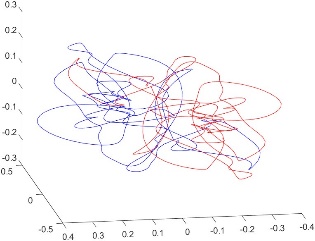

1000-500ms	500-0ms	1000-500ms	500-0ms

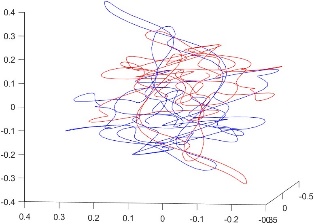

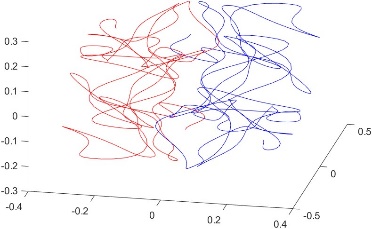

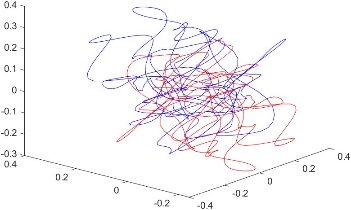

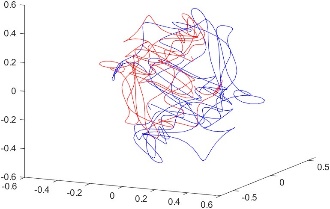

1000-500ms	500-0ms	1000-500ms	500-0ms

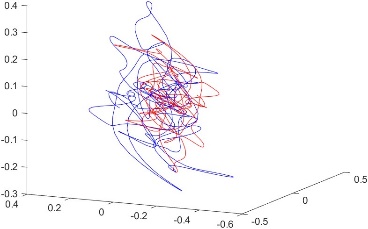

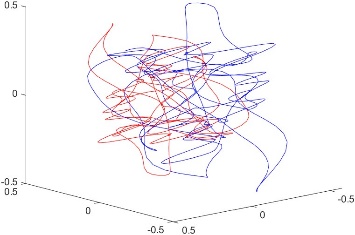

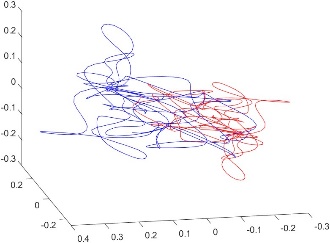

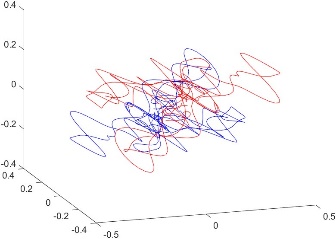

1000-500ms	500-0ms	1000-500ms	500-0ms

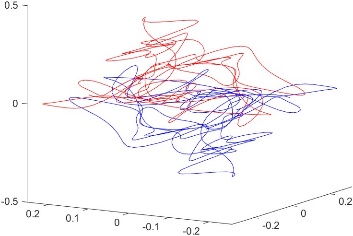

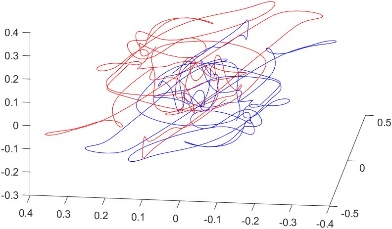

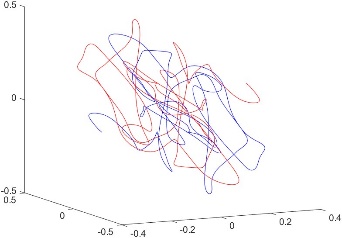

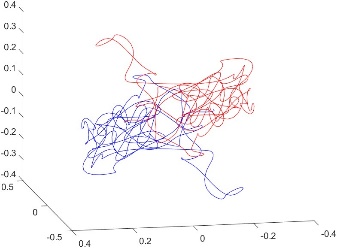

1000-500ms	500-0ms	1000-500ms	500-0ms

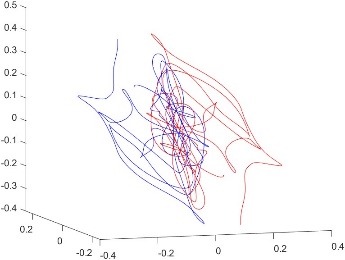

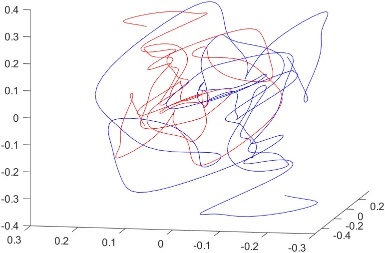

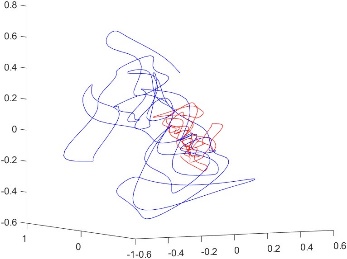

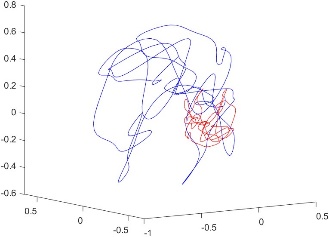

1000-500ms	500-0ms	1000-500ms	500-0ms
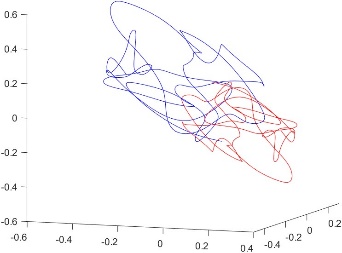

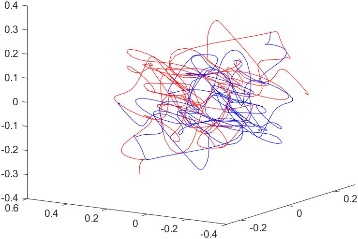

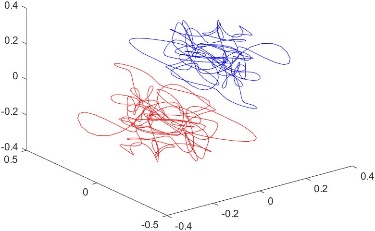

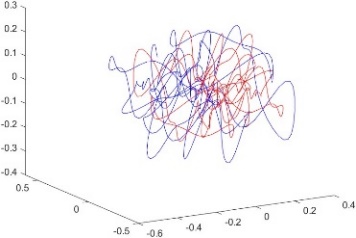

1000-500ms	500-0ms	1000-500ms	500-0ms

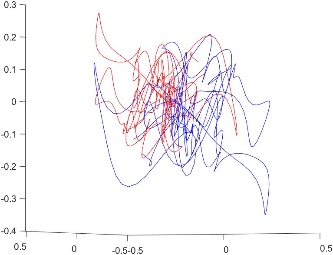

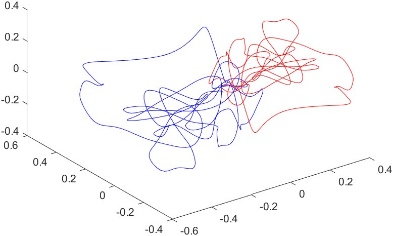

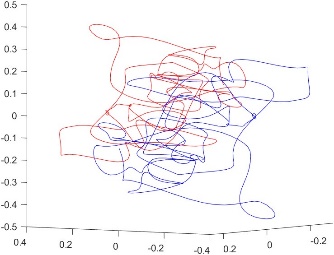

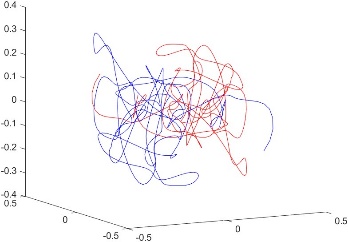

1000-500ms	500-0ms	1000-500ms	500-0ms

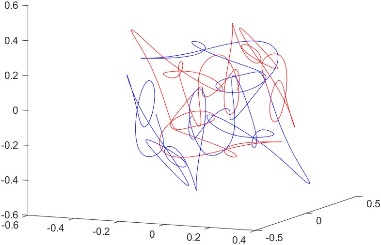

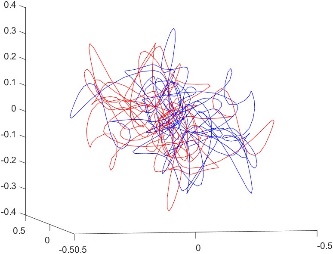

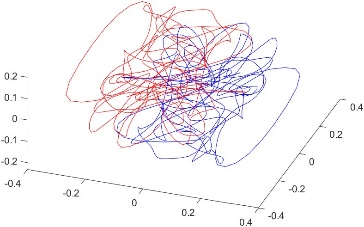

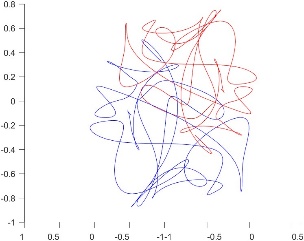

1000-500ms	500-0ms	1000-500ms	500-0ms

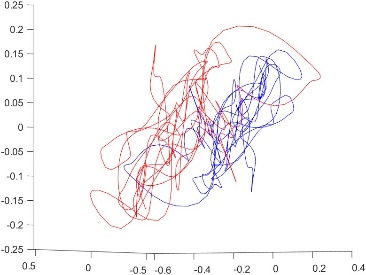

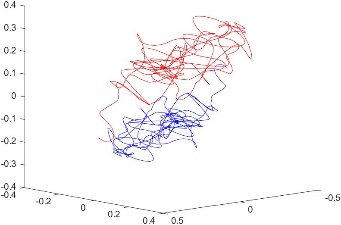

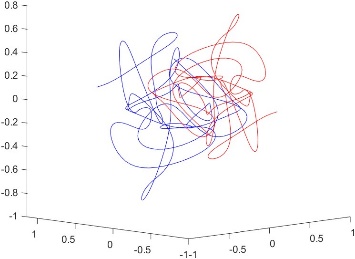

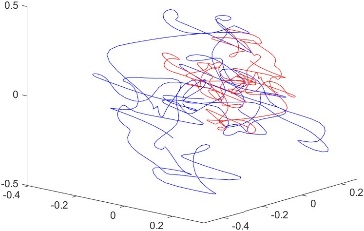

1000-500ms	500-0ms	1000-500ms	500-0ms

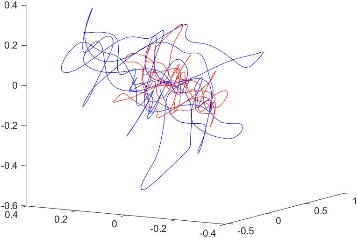

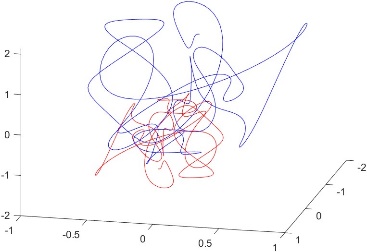

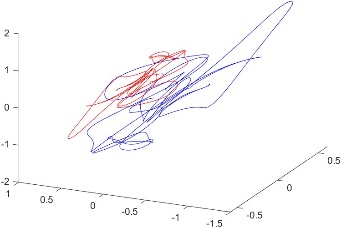

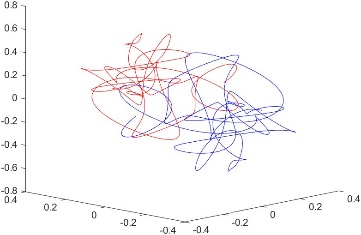

1000-500ms	500-0ms	1000-500ms	500-0ms

**Supplemental Data Figure 3.** The three latent variables that best separate the average safe and gamble latent variables graphed in 3-space. For each subject, the trajectories are graphed for the 1000-500ms and 500-0ms before choice. Red are gamble trials and blue are safe trials.
